## Supplementary material for "Evolutionary Adaptation of an HP1-protein Chromodomain Integrates Chromatin and DNA Sequence Signals": Key Resource Table

| **Key Resources Table** | | | | |
|---|---|---|---|---|
| **Reagent type (species) or resource** | **Designation** | **Source or reference** | **Identifiers** | **Additional information** |
| Antibody | anti-CG2678#2 (Rabbit polyclonal) | Baumgartner et al. 2022 | CG2678#2_4 P39glyc, raised against Kipferl peptide R171-I190 | Anti-Kipferl polyclonal antibody, available from Brennecke lab; ChIP (7 uL per IP) |
| Antibody | anti-CG2678 M3 (Mouse monoclonal) | Baumgartner et al. 2022 | M3 2C5-3C3, raised against Kipferl amino acids M2-K188 | Anti-Kipferl monoclonal IF antibody, available from Brennecke lab; IF (1:500) |
| Antibody | anti-Rhino (Rabbit polyclonal) | Mohn et al 2014 | Rhino#1_357 3gly | ChIP (5 uL per IP), IF (1:1000) |
| genetic reagent (D. melanogaster) | w1118;;; | Bloomington stock 3605 | w1118 | wildtype, cultivated in our lab for several years |
| genetic reagent (D. melanogaster) | w;; CG2678[Δ1](dsRed+)/TM3,Sb; | Baumgartner et al. 2022 | Kipferl (CG2678) | Kipferl mutant allele, available from VDRC; LB1-RMCEm31 |
| genetic reagent (D. melanogaster) | w;; CG2678[fs1]/TM 3,Sb; | Baumgartner et al. 2022 | Kipferl (CG2678) | Kipferl mutant allele, available from VDRC; LB1-FSm52; indel (-7); sequence CCTGCGTCCT GGCCGTGC------- TTTCCGGTTCA AGTGGCAAAG CGAGCAGAG |

| | | | | |
|---|---|---|---|---|
| genetic reagent (D. melanogaster) | w; rhi[18-7]/CyO; ; | Andersen *et al*., 2017, VDRC-ID 313488 | Rhino (CG10683) | mutant allele |
| genetic reagent (D. melanogaster) | w ; rhi[g2m11]/CyO ;; | Baumgartner et al. 2022 | Rhino (CG10683) | Rhino mutant allele, available from VDRC; indel -7 ; seq: ATGTCTCGCA ACCA-------cc- AATCTTGGTCT GGTCGATGCA CCGCCTAATG |
| genetic reagent (D. melanogaster) | w; rhi[G31D]/CyO; ; | this paper | Rhino (CG10683) | mutant allele changing glycine at position 31 to aspartic acid; m3-5 |
